# Tuba activates Cdc42 during neuronal polarization downstream of the small GTPase Rab8a

**DOI:** 10.1101/2019.12.16.876011

**Authors:** Pamela J. Urrutia, Felipe Bodaleo, Daniel A. Bórquez, Victoria Rozes-Salvador, Cristopher Villablanca, Cecilia Conde, Mitsunori Fukuda, Christian González-Billault

## Abstract

The acquisition of neuronal polarity is a complex molecular process that involves several different cellular mechanisms that need to be finely coordinated to define the somatodendritic and axonal compartments. Amongst such mechanisms, cytoskeleton and membrane dynamics control both the morphological transitions that define neuronal polarity acquisition as well as provide molecular determinants to specific sites in neurons at a defined time point. Small GTPases from the Rab and Rho families are well known molecular determinants of neuronal differentiation. However, during axon specification, a molecular link that couples proteins from these two families has yet to be identified. In this paper, we describe the role of Tuba, a Cdc42-specific guanine nucleotide-exchange factor (GEF), in neuronal polarity through a Rab8a-dependent mechanism. Rab8a or Tuba gain-of-function generates neurons with supernumerary axons whereas Rab8a or Tuba loss-of-function abrogated axon specification, phenocopying the well-established effect of Cdc42 on neuronal polarity. Neuronal polarization associated to Rab8a is also evidenced *in vivo*, since a dominant negative version of Rab8a severely impaired neuronal migration.

Remarkably, Rab8a activates Cdc42 in a Tuba-dependent manner, and dominant negative mutants of both GTPases reciprocally prevent the effect over polarity acquisition in the gain-of-function scenarios. Our results strongly suggest that a positive feedback loop linking Rab8a and Cdc42 activities via Tuba, is a primary event in neuronal polarization. In addition, we identified the GEF responsible for Cdc42 activation that is essential to specify axons in cultured neurons.

## INTRODUCTION

Neurons are highly-polarized cells, in which sprouting and elongation of neurites, and their subsequent development into axons and dendrites, are key events during early neuronal differentiation (Barnes and Polleux, 2009; Bradke and Dotti, 2000; Dotti et al., 1988). These cellular processes are heavily-dependent on changes in cytoskeletal dynamics and vectorially-directed membrane traffic which are controlled by the Rho and Rab families of small GTPases, respectively, thus providing the driving force for growth cone pathfinding, turning and neurite elongation (Arimura and Kaibuchi, 2007; Cheng and Poo, 2012; Dotti et al., 1988; Lalli, 2014). The small Rho GTPase Cdc42, has emerged as a critical regulator of neuronal polarity, allowing the activation of the PAR protein polarity complex, that is composed of an atypical protein kinase C (aPKC) isoform, and the scaffolding proteins Par3 and Par6. This complex accumulates at the tips of axons promoting axon specification via actin filament remodelling (Lalli, 2009; Nishimura et al., 2005; Shi et al., 2003). While it is widely-accepted that Cdc42 activation is required to drive axon specification, to date, the identity of the guanine nucleotide-exchange factor (GEF) responsible for such activation during the acquisition of neuronal polarity is still unknown.

Many other cell types are also highly-polarized, including epithelial cells. Their polarization mechanism has been extensively described, and shares several similarities with neuronal polarization (Bonifacino, 2014). During epithelial polarization, Tuba, a specific GEF for Cdc42, is essential to drive epithelial cell polarization (Bryant et al., 2010; Pichaud et al., 2019). Tuba, also named dynamin-binding protein (DNMBP), is a multidomain scaffold protein with GEF activity, that is highly-concentrated at neuronal synapses, where it interacts with dynamin to regulate the actin cytoskeleton (Salazar et al., 2003). Such a regulatory process involves the activation of Cdc42 and its binding with several other proteins, including several actin dynamic regulators such as N-WASP (neural Wiskott-Aldrich syndrome protein), WIPF3 (WAS/WASL-interacting protein family member 3), WAVE1 (WASP-family verprolin homologous protein 1), WIRE (WASP binding protein), CYFIP2 (cytoplasmic FMR1-interacting protein 2), NAP1 (nucleosome assembly protein 1), and Ena/VASP (enabled/vasodilator-stimulated phosphoprotein) (Cestra et al., 2005; Kovacs et al., 2011; Salazar et al., 2003). Moreover, Tuba plays a significant role in long term memory formation (Casoli et al., 2012). However, a role in neuronal polarization has not been reported to date.

On other hand, it has been recently discovered that exocytic (secretory) and endocytic pathways are critical for neuronal differentiation and axonal elongation (Villarroel-Campos et al., 2016a; Villarroel-Campos et al., 2014; Villarroel-Campos et al., 2016b). In the secretory pathway, the small Rab GTPase Rab8 regulates the transport of exocytic vesicles from the trans-Golgi network (TGN) to the plasma membrane (Stenmark, 2009). A deficiency in Rab8 impairs axon formation and decreases neurite outgrowth in embryonic hippocampal neurons, diminishing anterograde movement of vesicles that accumulate in the Golgi apparatus (Huber et al., 1995), whereas the overexpression of a constitutively active mutant of Rab8a increases axonal outgrowth in mice cortical neurons (Furusawa et al., 2017). In epithelial cell polarity, Rab8 controls apical Cdc42 activation through Tuba (Bryant et al., 2010). Therefore, here we evaluated whether Tuba is also the GEF that activates Cdc42 downstream of Rab8a during axon specification. We report that Tuba regulates axon specification by activation of Cdc42, through a positive feedback loop between Rab8a and Cdc42. Altogether, these findings suggest that Tuba may be one of the linkers that connects the regulation of cytoskeletal dynamics and the directed vesicular traffic during neuronal polarization.

## MATERIAL AND METHODS

### Primary culture

Neuronal hippocampal cultures were prepared as previously described (Kaech and Banker, 2006). Briefly, hippocampi were dissected from E18 Sprague-Dawley rat embryos or E17 C57BL/6 mice embryos and treated with 0.25% trypsin in HBSS (Invitrogen) for 25 min at 37°C. The tissue was mechanically-dissociated, and neurons were seeded onto sterilized glass coverslips coated with poly-L-lysine (1 mg/ml, Sigma-Aldrich) in DMEM (Invitrogen) containing 10% horse serum, 1% penicillin/streptomycin, and 1% glutamine (Invitrogen) for 1 h. The medium was then replaced with neurobasal medium containing 2% B27 (Invitrogen), 1% Glutamax (Invitrogen), and 1% penicillin/streptomycin (maintenance medium).

For nucleofection experiments, neurons in suspension were electroporated using the 4D-nucleofector system prior to plating following the manufacturer’s instructions (Lonza). Untagged constructs were co-nucleofected with a GFP-coding plasmid at a 4:1 ratio to ensure a 98% co-transfection rate, according to internal controls (data not shown).

### Cell culture

The neuroblastoma N1E-115 cell line was obtained from the American Type Culture Collection (ATCC, CRL-2263™). Cells were cultured in complete DMEM (Invitrogen) containing 5% heat-inactivated fetal bovine serum (FBS) and 1% penicillin/streptomycin (Invitrogen) at 37°C and 5% CO_2_. Cells were transfected in serum-free DMEM with Lipofectamine 2000 (Invitrogen) according to the manufacturer’s instructions. The culture medium was replaced with fresh medium at 4 h post-transfection and cells were processed 24–48 h later.

### Plasmids

The Raichu-Cdc42 Förster resonance energy transfer (FRET) biosensor probe was obtained from A. Cáceres (Instituto de Investigación Médica Mercedes y Martín Ferreyra, Córdoba, Argentina). Rab8aWT-GFP, Rab8aQ67L-GFP, Rab8aT22N-GFP, Rab8bWT-GFP, Rab8bQ67L-GFP, Rab8bT22N-GFP, Rab8aWT-Flag, Rab8aQ67L-Flag, Rab8aT22N-Flag, Rab8bQ67L-Flag, Rab8bQ67L-Flag, and Rab8bT22N-Flag (GFP and Flag are N-terminally tagged) plasmids were prepared as described previously (Fukuda, 2003; Homma and Fukuda, 2016). shRNAs against rat Rab8a cloned into the pRFP-C-RS vector (TF712324, Locus ID 117103), shRNAs against rat Rab8b cloned into the pGFP-V-RS vector (TG713207, Locus ID 266688) and shRNAs against mouse DNMBP cloned into the pRFP-C-RS vector (TF515449, Locus ID 71972) were purchased from OriGene. Cdc42F28L-HA was provided by Richard A. Cerione (Cornell University, Ithaca, New York, USA) and Cdc42-myc was purchased from Addgene.

### Antibodies

The following primary antibodies were used in this study: mouse anti-α-tubulin (clone DM1A, T6199; Sigma-Aldrich), rabbit anti-MAP2 (AB5622, Millipore), rabbit anti-Rab8 (D22D8, Cell Signaling), rabbit anti-Tuba (AB154836, Abcam), mouse anti-tau-1 (MAB3420, Millipore), mouse anti-βIII-tubulin (G712A; Promega), rabbit anti-Cdc42 (07-1466; Millipore), mouse anti-cofilin and rabbit anti-phospho-cofilin (Ser3) (a kind gift from J. R. Bamburg, Colorado State University, Colorado, USA). Secondary antibodies for immunoblots were HRP-conjugated anti-mouse and anti-rabbit IgG (Jackson Laboratories). Secondary antibodies for immunocytochemistry were anti-mouse and anti-rabbit conjugated to Alexa-Fluor 488, 543 or 633 (Thermo Fischer).

### Cdc42 activity pull-down assay

The CRIB domain for Cdc42 activity was purified as described previously (Villarroel-Campos et. al, 2016b). Briefly, loaded beads were incubated for 70 min at 4°C with 1.5 mg of either a control or Cdc42-expressing N1E-115 cell lysate using fishing buffer (50 mM Tris-HCl, pH 7.5, 10% glycerol, 1% Triton X-100, 200 mM NaCl, 10 mM MgCl_2_, 25 mM NaF, and protease inhibitor cocktail). The beads were washed three times with washing buffer (50 mM Tris-HCl, pH 7.5, 30 mM MgCl_2_, 40 mM NaCl), and then resuspended in SDS-PAGE sample buffer. Bound Cdc42-GTP was subjected to immunoblot analysis and quantified with respect to total Cdc42 using ImageJ.

### Cdc42 activity FRET assay

To measure GTPase activity, neurons were co-transfected with Flag-Rab8a (Q67L or T22N) and Raichu-Cdc42 FRET biosensor for 24 h. FRET efficiency measurements were performed as described previously (Nakamura et al., 2006). Briefly, transfected neurons were excited at 450 nm, and emissions were collected at 460–490 and 505–530 nm (donor and acceptor emission wavelengths, respectively). The ratio of the acceptor-to-donor emission was established as the FRET efficiency. The FRET map was generated by dividing the acceptor-to-donor ratio image by the binary mask of the same image. Measurement of FRET efficiency was performed by selecting a region of interest at the soma, and at the proximal or distal axon.

### Immunoblotting

Protein extracts were obtained from cell lines or primary neuronal cultures. Cells were lysed with radioimmunoprecipitation assay (RIPA) buffer, subjected to SDS-PAGE and electroblotted onto nitrocellulose or polyvinylidene difluoride membranes. Blots were probed with the appropriate primary and secondary antibodies, and immunoreactivity signals were visualized with an enhanced chemiluminescent substrate (Thermo Scientific) and quantified by densitometry using ImageJ.

### Immunocytochemistry

Cultures were fixed with 4% PFA/4% sucrose in PBS for 30 min at 37°C, washed three times with PBS and then permeabilized with 0.2% Triton X-100 in PBS. Then coverslips were blocked with 5% BSA in PBS, primary antibodies were added in 1% BSA in PBS, and cells were incubated overnight at 4°C. Cells were washed three times with PBS and incubated with the appropriate AlexaFluor-conjugated secondary antibodies for 1h at room temperature. Coverslips were mounted in FluorSave (Millipore) and analyzed using a confocal microscope (Zeiss LSM 810). Axon length was measured using the microscope-associated software, LSM Image Browser (Zeiss). Confocal images presented here were color-inverted to improve morphological appreciation.

### In utero electroporation (IUE) and imaging acquisition

IUE were carried out following previous reports (Fuentes et al., 2012; Mestres et al., 2016). Briefly, pregnant E15.5 C57BL/6 mice were anesthetized with isofluorane/oxygen mix (4% for induction and 2% for maintenance) during the whole surgery, and Tramadol (5 mg/Kg) was used as analgesia during the procedure. Uterine horns were exposed and embryos were locally-injected with pCAG-GFP, pCAG-GFP+Rab8aQ67L-GFP or pCAG-GFP+Rab8aT22N-GFP into the lateral ventricle of the brain. To visualize successful injections, the fast green FCF dye (Sigma-Aldrich, catalog number: F7252) was co-injected with DNAs. Then, brains were electroporated using a BTX electroporator (ΔV= 39 V; pulses: 5; duration: 50 ms; intervals between pulses: 950 ms) with Tweezers w/Variable Gap 2 Square Platinum Electrodes (Nepagene, CUY647P2X2). After electroporation, in utero embryos were returned to the maternal cavity to recover from the surgery. At E18, embryos were sacrificed to check GFP expression in control and Rab8aT22N or Rab8aQ67L genetic contexts. Brains expressing GFP were fixed overnight in 4% w/v PFA solution dissolved in PBS at 4°C with gentle agitation. Then, fixed brains were immersed into a 30% w/v sucrose solution for 24 h at 4°C. Post-fixed brains were frozen at −25°C using Crioplast solution (Biopack). The cerebral cortex was sliced into 50 µm cortical sections using a cryostat (Leica CM 1850). Brain slices were mounted onto glass slides. Tissues were permeabilized with 0.3% v/v Triton X-100-PBS solution, followed by DAPI staining (15 min at RT). Then, samples were mounted in Mowiol solution (Sigma) for z-stack imaging in a Zeiss LSM 810 confocal microscopy. Images were acquired with a 20x air objective. Several fields were acquired in order to image the whole brain cortex (from the ventricular zone to the cortical plate). Collected images were stitched using the Stitching plug-in of Fiji-Image J.

### Statistical analyses

All data represent mean□±□SEM of at least three independent experiments. Comparisons between two groups were made using unpaired Student’s t-tests when data presented a Gaussian distribution, and non-parametric Mann-Whitney tests when data presented a non-Gaussian distribution. Comparisons between more than two groups were carried out using one-way ANOVA followed by Bonferroni’s post-test. A value of p□<□0.05 was considered significant.

## RESULTS

### Tuba, a Cdc42 GEF, is required for axon specification during neuronal polarization

In light of its role in epithelial cell polarity, we first analyzed the expression, distribution and function of Tuba during neuronal differentiation. Tuba is expressed in adult brain and co-localizes with synaptic markers (Salazar et al., 2003). Two isoforms of Tuba have been described, Tuba full-length (TubaFL) (∼180 kDa) and miniTuba (mTuba) (∼75 kDa), differing by the inclusion (by alternative splicing) of four N-terminal SH3 (Src-homology 3) domains (Salazar et al., 2003). We detected increasing levels of TubaFL (using an isoform-specific antibody) during neuronal polarization in cultured neurons at several days *in vitro* (div)(Figure 1A). Immunofluorescence staining of neurons showed a punctate pattern with perinuclear distribution in stage 1 neurons. In unpolarized stage 2 neurons, Tuba accumulated in a single minor neurite and upon polarization (Stage 3) it became enriched in the axonal hillock and axonal growth cone (Figure 1B). To explore whether Tuba is involved in axon formation, we performed knockdown experiments in cultured hippocampal neurons using an shRNA against Tuba. The efficiency of the shRNA construct was tested by western blot in N1E-115 neuroblastoma cells (Supplementary Figure 1). Mouse hippocampal neurons were nucleofected before plating with an shRNA-encoding vector and analyzed at 3 div by immunofluorescence staining against MAP2 and tau1, as dendritic and axonal markers, respectively (Figure 1C). Tuba knockdown increased the number of unpolarized neurons (41%) compared to scrambled shRNA-transfected neurons (Figure 1D). In addition, we analyzed the minor neurite mean length and total axonal length of nucleofected neurons. Tuba knockdown did not alter minor neurite mean length (Figure 1E) although it did generate a significant decrease in axonal length as compared to control neurons (Figure 1F). These results indicate that Tuba is required for axon formation. In order to confirm the role of Tuba in neuronal polarization, we nucleofected before plating hippocampal neurons with TubaFL, mTuba or mTuba without the GEF domain (mTubaΔGEF) constructs and analyzed them at 3 div using immunofluorescence staining against MAP2 and tau-1 (Figure 1G). TubaFL nucleofection increased the number of neurons showing multiple axons (38%), whereas mTuba did not. mTubaΔGEF nucleofection increased the number of unpolarized neurons (40.6%) similar to what was observed in the Tuba knockdown experiments (Figure 1H). We also found an increase in mean minor neurite length in neurons transfected with mTuba; however we observed no significant differences after TubaFL or mTubaΔGEF nucleofection (Figure 1I). Furthermore, we detected a reduction in the total axonal length in mTubaΔGEF nucleofected neurons compared to control, but no differences in the total axonal length between TubaFL or mTuba expressing neurons (Figure 1J), suggesting that the role of Tuba in axon formation is partially-dependent of its GEF activity.

**Figure 1.**
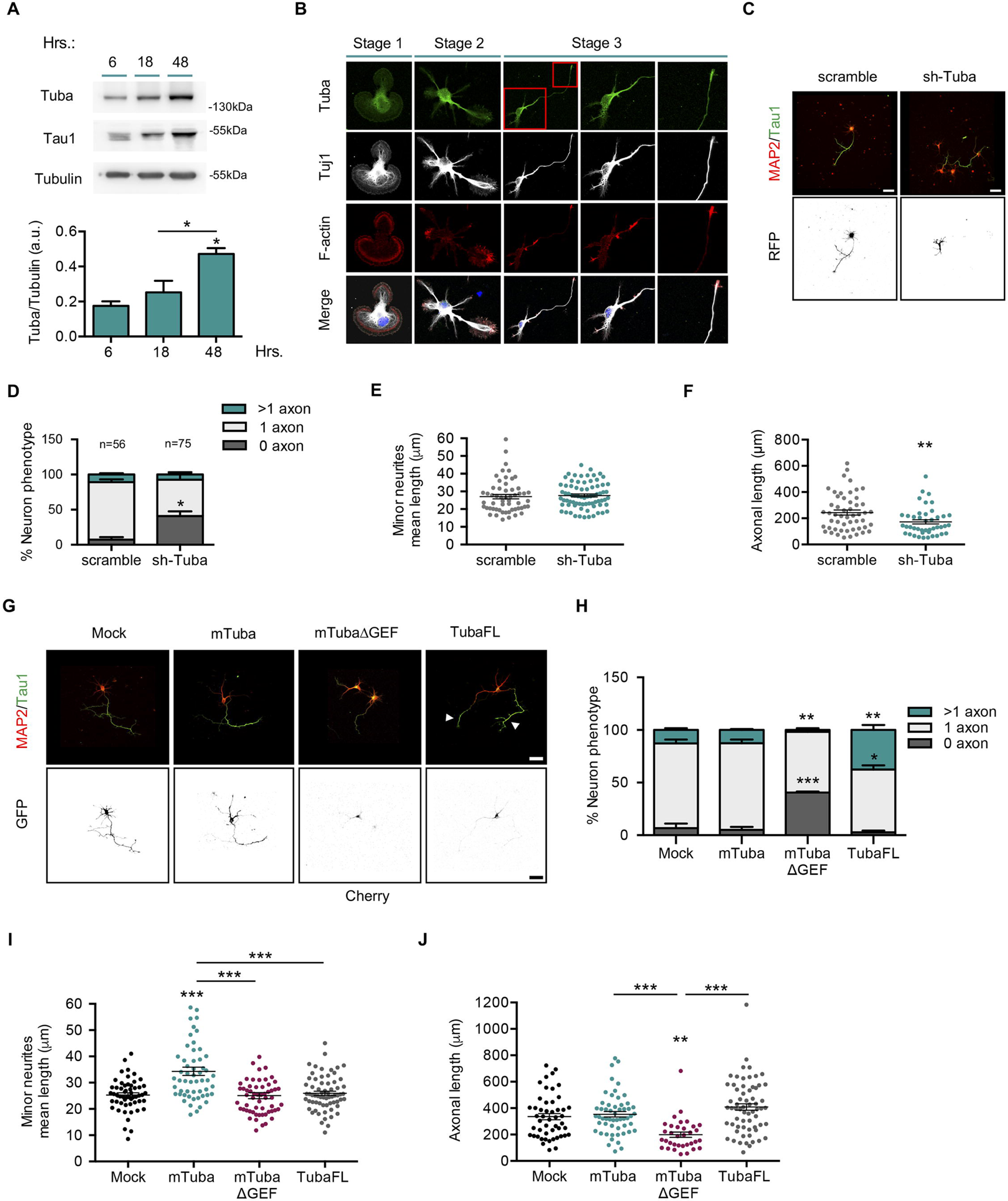
Tuba, a Cdc42 GEF, is required for axon specification during neuronal polarization. (**a**) Representative western blot and quantification of Tuba expression from cultured cortical neurons at 6, 18 and 48 h post-plating. (**b**) Representative images of neurons at stage 1, 2 and 3 stained with Tuba (green), Tuj1 (grey), F-actin (red) and TO-PRO-3 (blue). **(c)** Representative images of hippocampal neurons overexpressing scrambled or shRNA against Tuba, stained with MAP2 and tau-1. (**d**) Quantification of neuronal phenotypes of RFP positive cells in (c). (**e**) Quantification of minor neurite average length in (c). (**f**) Quantification of axonal length in (c). (**g**) Representative images of hippocampal neurons overexpressing mock, mini Tuba (mTuba), mtubaΔGEF or Tuba Full-length (TubaFL), stained with MAP-2 and tau-1. Arrowheads show axons in multitaxonic neurons. (**h**) Quantification of neuron phenotypes of GFP or Cherry positive cells in (g). (**i**) Quantification of mean minor neurite length in (g). (**j**) Quantification of axonal length in (g). Scale bars, 50 μm. Values represent mean ± SEM (n = 3 or 4). *p<0.05, **p<0.01 and ***p<0.001 as compared to the corresponding control.

### Tuba is associated with Rab8a at the Golgi compartment

As only TubaFL (and not mTuba) can interact with GM130, a peripheral membrane protein of the cis-Golgi stack, that controls Cdc42 activation at the Golgi apparatus (Kodani et al., 2009), and Rab8 was originally-described as a trafficking regulator between the Golgi apparatus and the plasma membrane, we then evaluated the co-distribution of Tuba and Rab8 with GM130. We performed immunofluorescence staining at 3 div in hippocampal neurons overexpressing TubaFL-GFP and immunostained against endogenous Rab8 and GM130. We observed that Rab8 and Tuba partially co-localized with the GM130-positive compartment and also at the axonal growth cone (Figure 2A), with Pearsońs correlation coefficient of 0.28 ± 0.07 and 0.17 ± 0.044, respectively (Figure 2B).

**Figure 2.**
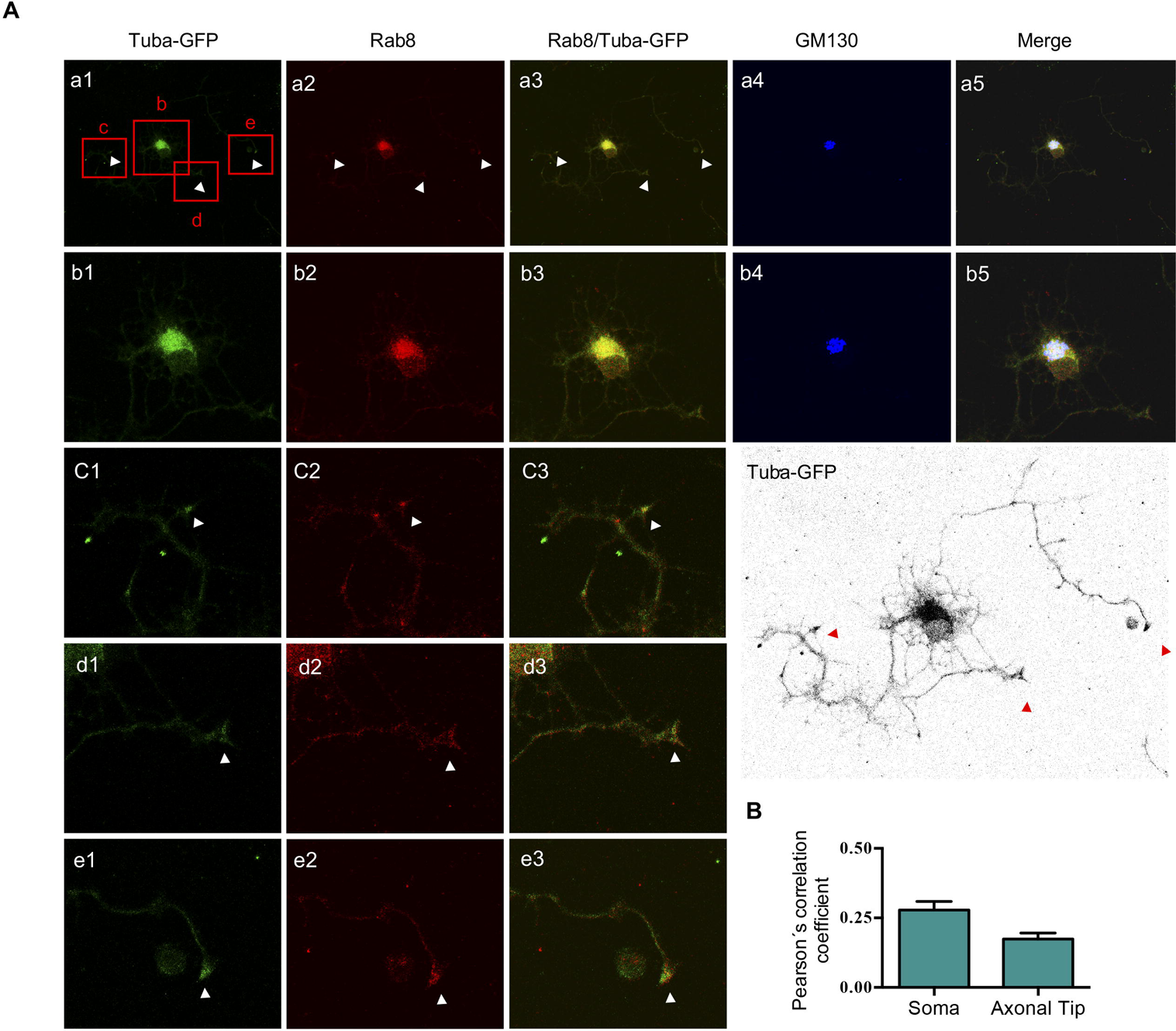
Tuba is associated with Rab8a at the Golgi compartment. (**a**) Representative image of Tuba-transfected neurons stained with anti-rab8 (red) and anti-GM130 (cis-Golgi marker, blue) antibodies. Columns 1, 2, 3 and 4 correspond to Tuba-GFP, Rab8, Rab8/Tuba-GFP, GM130 and merge, respectively. Rows b, c, d and e correspond to a magnification indicated in row a. A color-inverted TubaFL overexpressing neuron is shown on the lower right side of the panel. (**b**) Quantification of Pearson’s correlation coefficient in soma at neuronal tips of (a).

### Rab8 is required for axon formation

To examine whether Rab8 is required for axon specification, we analyzed the expression, distribution and function of Rab8 during early neuronal differentiation. In cultured neurons, we detected the expression of Rab8 (∼25 kDa) as early as 6 h *in vitro*, expression which increased during neuronal polarization (Figure 3A). Staining of cultured neurons showed a punctate pattern with perinuclear distribution, present in all minor neurites in unpolarized cells (Stage 1 and 2) and in axons and minor neurites of polarized cells (Stage 3) (Figure 3B). To explore whether Rab8 was involved in axon formation, we performed knockdown experiments in cultured hippocampal neurons using an shRNA against Rab8. There are two Rab8 proteins, termed Rab8a and Rab8b, which are encoded by different genes (Armstrong et al., 1996); therefore we used isoform-specific shRNAs to distinguish their individual contribution to axonal development. The efficiency of each shRNA was assessed by western blot in transfected-B104 neuroblastoma cells (Supplementary Figure 2). Hippocampal neurons were then nucleofected before plating with shRNA against Rab8a and/or Rab8b, and analyzed at 3 div using immunofluorescence staining against MAP2 and tau-1 (Figure 3C, 3F and 3K). Rab8a knockdown increased the number of unpolarized neurons (37.3%) compared to scrambled (Figure 3D). Additionally, we analyzed the mean minor neurite length and total axonal length of nucleofected neurons at 3 div. Rab8a knockdown did not alter mean minor neurite length (Figure 3E) but lead to a reduction in total axonal length (Figure 3F). In contrast, Rab8b knockdown did not alter the number of unpolarized neurons or the mean minor neurite length (Figure 3H-I), but reduced total axonal length (Figure 3J). In addition, co-nucleofection using both shRNAs against Rab8a and Rab8b, caused an increase in the number of unpolarized neurons (Figure 3L) that was similar to what we observed with shRab8a. Moreover, Rab8a and the double knockdown did not alter mean minor neurite length (Figure 3M) but generated a reduction in total axonal length (Figure 3N). Collectively, these results argue that Rab8a, but not Rab8b, could be required for axon formation.

**Figure 3.**
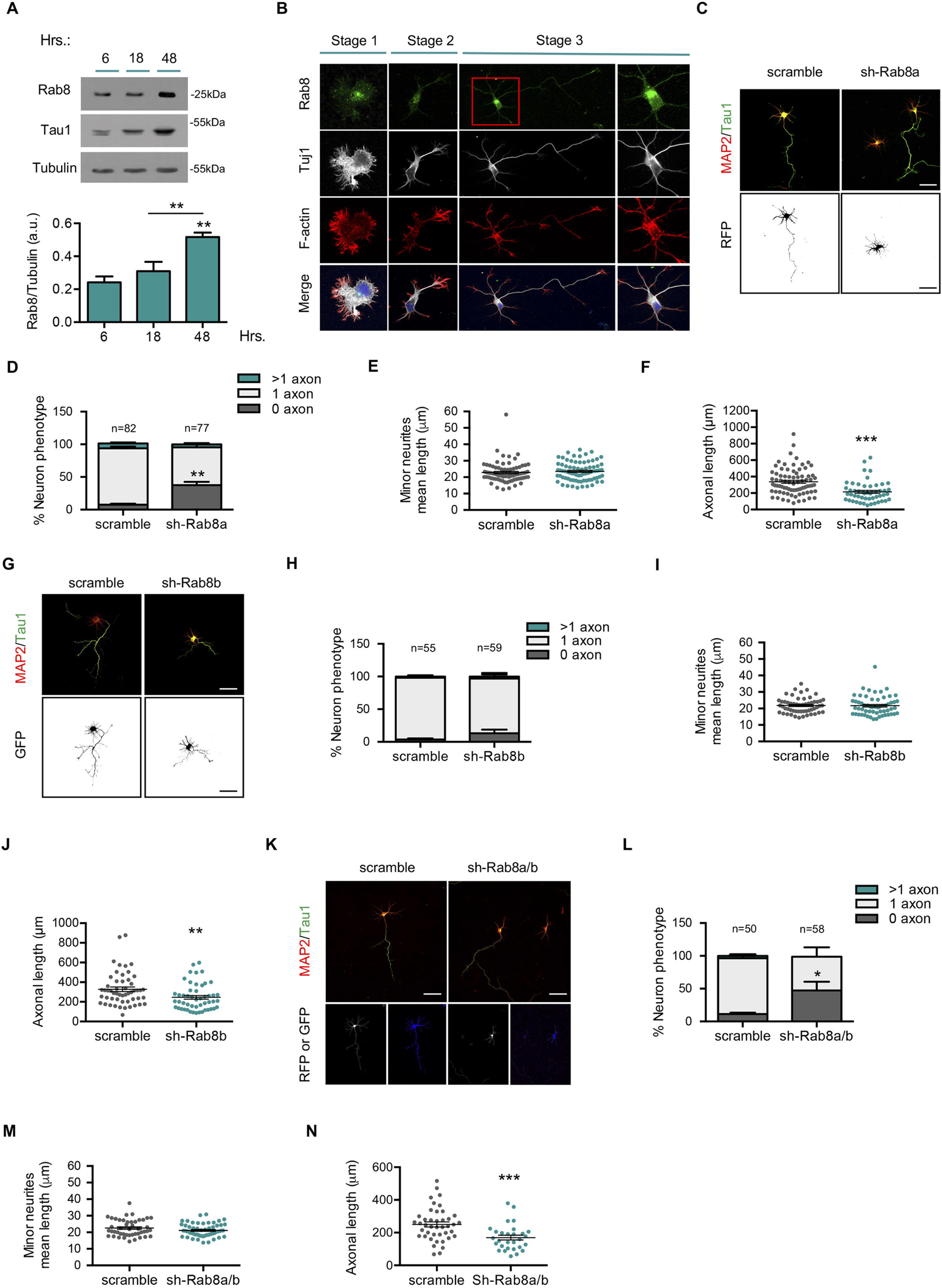
Rab8a is required for axon formation. (a) Representative western blot and quantification of Rab8 expression from cultured cortical neurons at 6, 18 and 48 h post-plating. (**b**) Representative images of neurons at stage 1, 2 and 3 stained with Rab8 (green), Tuj1 (grey), F-actin (red) and TO-PRO-3 (blue). **(c)** Representative images of hippocampal neurons overexpressing scrambled or shRNA against Rab8a, stained with MAP2 and tau-1. (**d**) Quantification of neuron phenotypes of RFP positive cells in (c). (**f**) Quantification of mean minor neurite length in (c). (**f**) Quantification of axonal length in (c). (**g**) Representative images of hippocampal neurons overexpressing scrambled or shRNA against Rab8b, stained with MAP2 and tau-1. (**h**) Quantification of neuron phenotypes of GFP positive cells in (g). (**i**) Quantification of mean minor neurite length in (g). (**j**) Quantification of axonal length in (g). (**k**) Representative images of hippocampal neurons overexpressing scrambled or shRNA against Rab8a and Rab8b (1:1), stained with MAP2 and tau-1. (**l**) Quantification of neuron phenotypes of GFP and RFP positive cells in (k). (**m**) Quantification of mean minor neurite length in (k). (**n**) Quantification of axonal length in (k). Scale bars, 50 μm. Values represent mean ± SEM (n=3). *p<0.05, **p<0.01 and ***p<0.001 as compared to the corresponding control.

### Overexpression of a constitutively active (CA) Rab8a isoform generates multiaxonal neurons

To further analyze the role of Rab8a in axon specification, we nucleofected Rab8a and Rab8b WT, Rab8a and Rab8b Q67L (constitutively active, CA mutants), and Rab8a and Rab8b T22N (dominant negative, DN mutants) as eGFP fusion proteins in cultured hippocampal neurons and addressed polarity acquisition by staining against MAP2 and tau-1 at 3 div (Figure 4A, 4E). The overexpression of Rab8a WT and Rab8a Q67L lead to a significant increase in the number of neurons showing multiple axons (30% and 55% more, respectively), whereas the overexpression of Rab8a T22N caused a significant increase in the number of unpolarized neurons (50% more) compared to control conditions (Figure 4B). In addition, we measured other morphometric parameters. Neurons overexpressing Rab8a Q67L showed no differences in the minor neurite length but had significant longer axons compared to the control. Neurons overexpressing Rab8a T22N displayed a significant decrease in both mean minor neurite length and axonal length compared to the control (Figure 4C-D). To unequivocally determine that multiple tau-1 positive protrusions were indeed axons, we performed immunofluorescence against ankyrin G, a marker of the axonal initial segment (AIS), observing that neurons overexpressing Rab8a Q67L have several AISs (Supplementary Figure 3). In contrast, the overexpression of Rab8b WT or Rab8b Q67L generated no variations in the number of neurons showing multiple axons, although the overexpression of Rab8b T22N resulted in a significant increase in the number of unpolarized neurons (Figure 4F). We did not observe differences in mean minor neurite length in neurons overexpressing Rab8b WT, Rab8b Q67L or Rab8b T22N compared to the control (Figure 4G). However, neurons overexpressing Rab8b T22N had significantly shorter axons compared to the control neurons (Figure 4H). These results clearly indicate that Rab8a is important for axon specification, because its absence or the expression of its inactive form abolished axon generation, whereas the overexpression of a Rab8 WT or a constitutively active mutant induced the generation of supernumerary axons.

**Figure 4.**
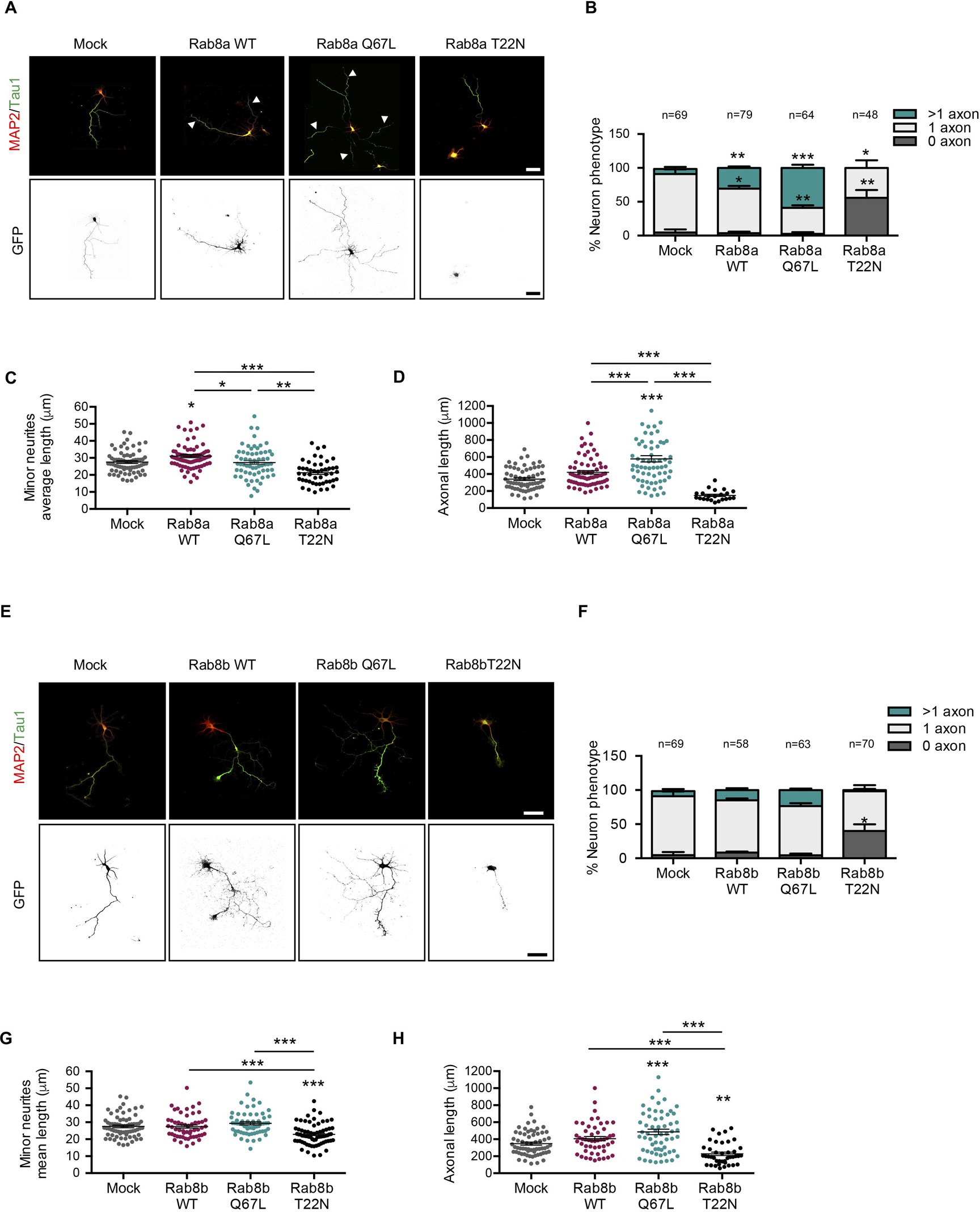
Overexpression of a constitutively active (CA) Rab8a isoform generates multiaxonic neurons. (**a**) Representative images of hippocampal neurons overexpressing Rab8a WT, Rab8a Q67L (CA mutant) or Rab8a T22N (dominant negative, DN, mutant), stained with MAP-2 and tau-1. Arrowheads indicate axons in the multiaxonic neurons. (**b**) Quantification of neuron phenotypes of GFP positive cells in (a). (**c**) Quantification of mean minor average length in (a). (**d**) Quantification of axonal length in (a). (**e**) Representative images of hippocampal neurons overexpressing Rab8b WT, Rab8b Q67L (CA mutant) or Rab8b T22N (DN mutant), stained with MAP2 and tau-1. (**f**) Quantification of neuron phenotypes of GFP positive cells in (e). (**g**) Quantification of minor neurite average length in (e). (**h**) Quantification of axonal length in (e). Scale bars, 50 μm. Values represent mean ± SEM (n=3). *p<0.05, **p<0.01 and ***p<0.001 as compared to the corresponding control.

### Rab8a CA triggers Tuba-dependent Cdc42 activation

It has been previously-reported that Rab8 controls apical differentiation in epithelial cells by Cdc42 activation through recruitment of Tuba (Bryant et al., 2010).

Therefore, we next evaluated the effect of Rab8a activation on Cdc42 activity and the phosphorylation of Serine 3 of cofilin, a downstream indicator of Cdc42 signalling pathway activation. Neuroblastoma N1E-115 cells were transfected with Rab8a WT, Rab8a Q67L or Rab8a T22N and Cdc42 activity was assessed by a pulldown assay. The overexpression of Rab8a Q67L induced a significant increase in Cdc42 activity and phospho-cofilin compared with the control (Figure 5A and B). We also evaluated changes in Cdc42 activity in cultured primary neurons using a FRET-based approach. We expressed a Cdc42 FRET biosensor in conjunction with Rab8a Q67L or Rab8a T22N and measured pixel intensity in the resulting FRET maps in the somatodendritic, proximal axon or distal axon compartments (Figure 5C). Neurons overexpressing Rab8a Q67L had significantly more Cdc42 activity in all compartments compared to the control. No changes in Cdc42 activity were observed in neurons overexpressing Rab8a WT or Rab8a T22N (Figure 5D-E). In addition, we evaluated the effect of the knockdown of Tuba on Cdc42 activation induced by Rab8a Q67L in a neuroblastoma cell line. We observed that shRNA against Tuba abrogated the increased Cdc42 activity induced by Rab8a Q67L (Figure 5G). We also analyzed the effect of shRNA against Tuba in the neuronal morphological changes generated by Rab8a Q67L (Figure 5H). We co-nucleofected an shRNA against Tuba and Rab8a Q67L in primary cultures of hippocampal neurons, and observed that shRNA against Tuba reversed the generation of supernumerary axons induced by Rab8a Q67L, and increased the number of neurons without axons (Figure 5I). No changes were observed in the mean minor neurite length, but significant reductions in axonal length in the neurons overexpressing Rab8a Q67L and shRNA against Tuba compared with Rab8a Q67L alone (Figure 5J-K) were detected. Therefore, these results suggest that increased Cdc42 activity induced by Rab8a, could be mediated by Tuba and that the generation of neurons showing multiple axons mediated by Rab8 overexpression could also be dependent on Tuba expression.

**Figure 5.**
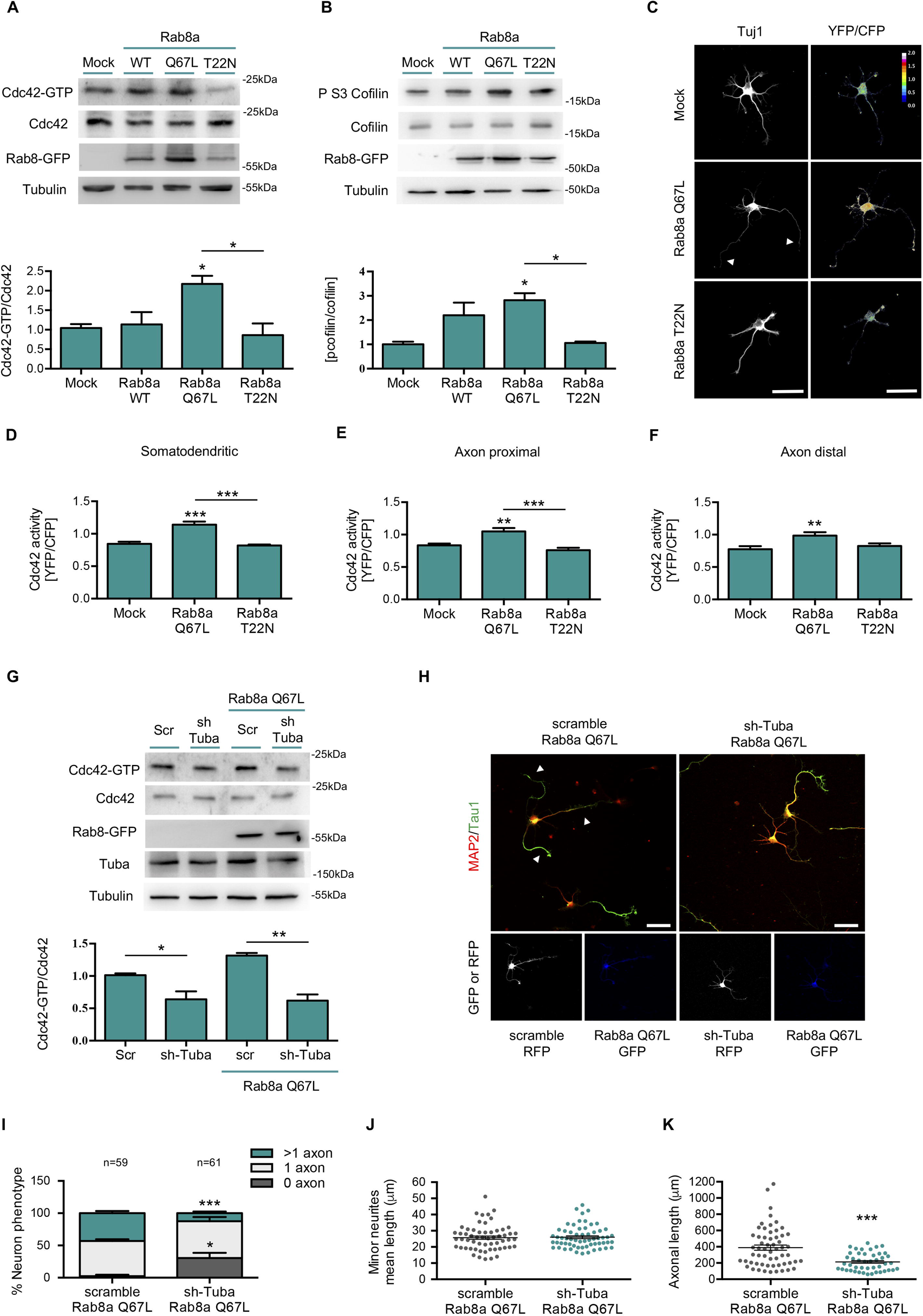
Rab8a CA triggers Tuba-dependent Cdc42 activation. (**a**) Representative image and quantification of pulldown assays from N1E-115 neuroblastoma cell lines overexpressing GFP, Rab8a WT, Rab8a Q67L (CA mutant) or Rab8a T22N (DN mutant). (**b**) Representative image and quantification of western blots for cofilin and phosphor ser3 cofilin from N1E-115 neuroblastoma cell lines overexpressing GFP, Rab8a WT, Rab8a Q67L (CA mutant) or Rab8a T22N (DN mutant). (**c**) 1 DIV cultured hippocampal neurons expressing a FRET biosensor for Cdc42 co-expressing Flag, Flag-Rab8a Q67L or Flag-Rab8a T22N and stained against Tuj1 (white). (**d**,**e**,**f**) Quantification of the Cdc42-GTP/Cdc42 ratio from (c) in somatodendritic (d), proximal axonal (e) and distal axonal compartments (f). (**g**) Representative image and quantification of pulldown assays from N1E-115 neuroblastoma cell lines knockdown for Tuba overexpressing GFP or Rab8a Q67L (CA mutant). (**h**) Representative images of Rab8a Q67L-overexpressing neurons co-transfected with a shRNA against Tuba, stained with MAP-2 and tau-1. Arrowheads show axons in multitaxonic neurons. (**i**) Quantification of neuron phenotypes of GFP and RFP positive cells in (h). (**j**) Quantification of minor neurite average length in (h). (**k**) Quantification of axonal length in (h). Scale bars, 50 μm. Values represent mean ± SEM (n=3). *p<0.05, **p<0.01 and ***p<0.001 as compared to the corresponding control.

### A positive feedback loop between Rab8a and Cdc42 regulates neuronal polarization

In the next set of experiments, we examined whether the overexpression of a Cdc42 fast cycling mutant reversed the loss of polarity induced by overexpression of Rab8a T22N in neurons. We performed co-nucleofection of different versions of Cdc42 and Rab8a and analyzed their effect on neuronal morphology by immunofluorescence against tau-1 and MAP-2 (Figure 6A). We observed that co-expression of Rab8a T22N reversed the generation of neurons showing multiple axons induced by Cdc42 F28L (fast cycling mutant), while the overexpression of Cdc42 T17N (dominant negative, DN mutant) reversed the generation of supernumerary axons induced by Rab8a Q67L. In addition, the concurrent overexpression of Cdc42 F28L and Rab8a Q67L generated a robust increase in the number of multiaxonal neurons, similar to that observed for each one separately (Figure 6B). We also discovered that co-expression of Rab8a T22N reduced the minor neurite length of Cdc42 F28L-overexpressing neurons and in turn, the co-expression of Rab8a Q67L increased the minor neurite length of Cdc42 T17N-overexpressing neurons (Figure 6C). Similarly, co-expression of Rab8a T22N reduced the axonal length of Cdc42 F28L-overexpressing neurons; however, we observed no differences with the co-expression of Rab8a Q67L in Cdc42 T17N-overexpressing neurons (Figure 6D). These results suggest the existence of a positive feedback loop between Rab8a and Cdc42 and that both activities are needed to regulate neuronal polarization.

**Figure 6.**
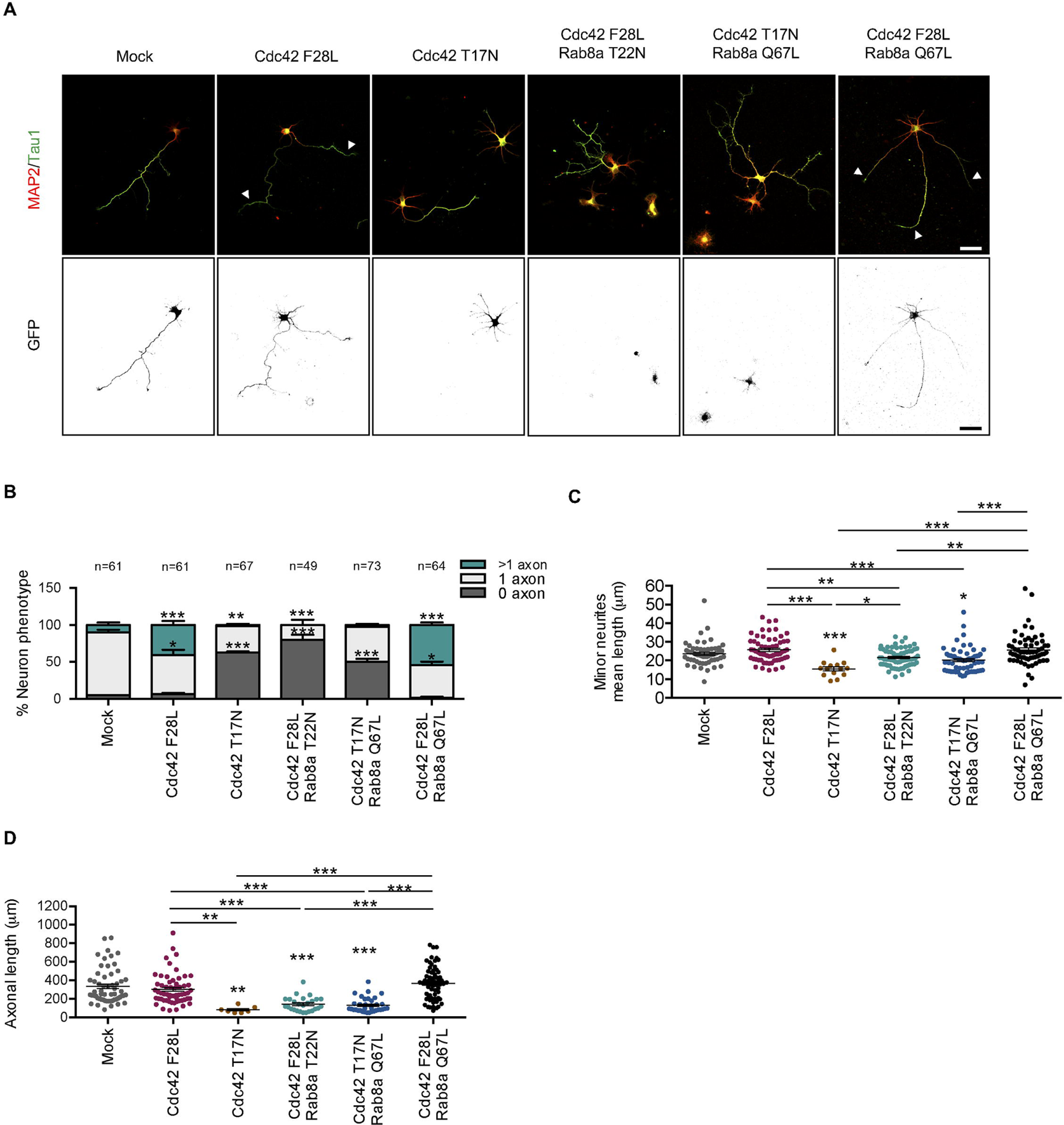
A positive feedback loop between Rab8a and Cdc42 regulates neuronal polarization. (**a**) Representative images of Cdc42 F28L (fast cycling mutant) or Cdc42 T22N (DN mutant)-overexpressing neurons co-transfected with Rab8a Q67L or Rab8a T22N, stained with MAP-2 and tau-1. Arrowheads show axons in multitaxonic neurons. (**b**) Quantification of neuron phenotypes of GFP positive cells in (a). (**c**) Quantification of minor neurite average length in (a). (**d**) Quantification of axonal length in (a). Scale bars, 50 μm. Values represent mean ± SEM (n=3). *p<0.05, **p<0.01 and ***p<0.001 as compared to the corresponding control.

### Rab8a regulates the neuronal morphology of migrating neurons at the embryonic cortex

In the final set of experiments, we sought to determine whether Rab8a could regulate neuronal polarization *in vivo*. To this end, we performed IUE in E15.5 embryos to express a pCAG-GFP vector with the cDNAs of Rab8a T22N or Rab8a Q67L into the lateral ventricle, followed by fixation at E18. We analyzed neuronal morphology in 4 different cortical layers: ventricular zone (VZ), subventricular zone (SVZ), intermediate zone (IZ) and cortical plate (CP) (Fuentes et al., 2012). Control electroporated neurons (visualized through eGFP) showed normal migration patterns. In contrast, the number of neurons expressing Rab8a T22N that reached the cortical plate was significantly reduced, compared to neurons overexpressing Rab8a Q67L (Figure 7A, C). In addition, we observed a notorious trend showing an increase in the number of neurons that remained at VZ and SVZ regions when Rab8a T22N was expressed (Figure 7A, C).

**Figure 7.**
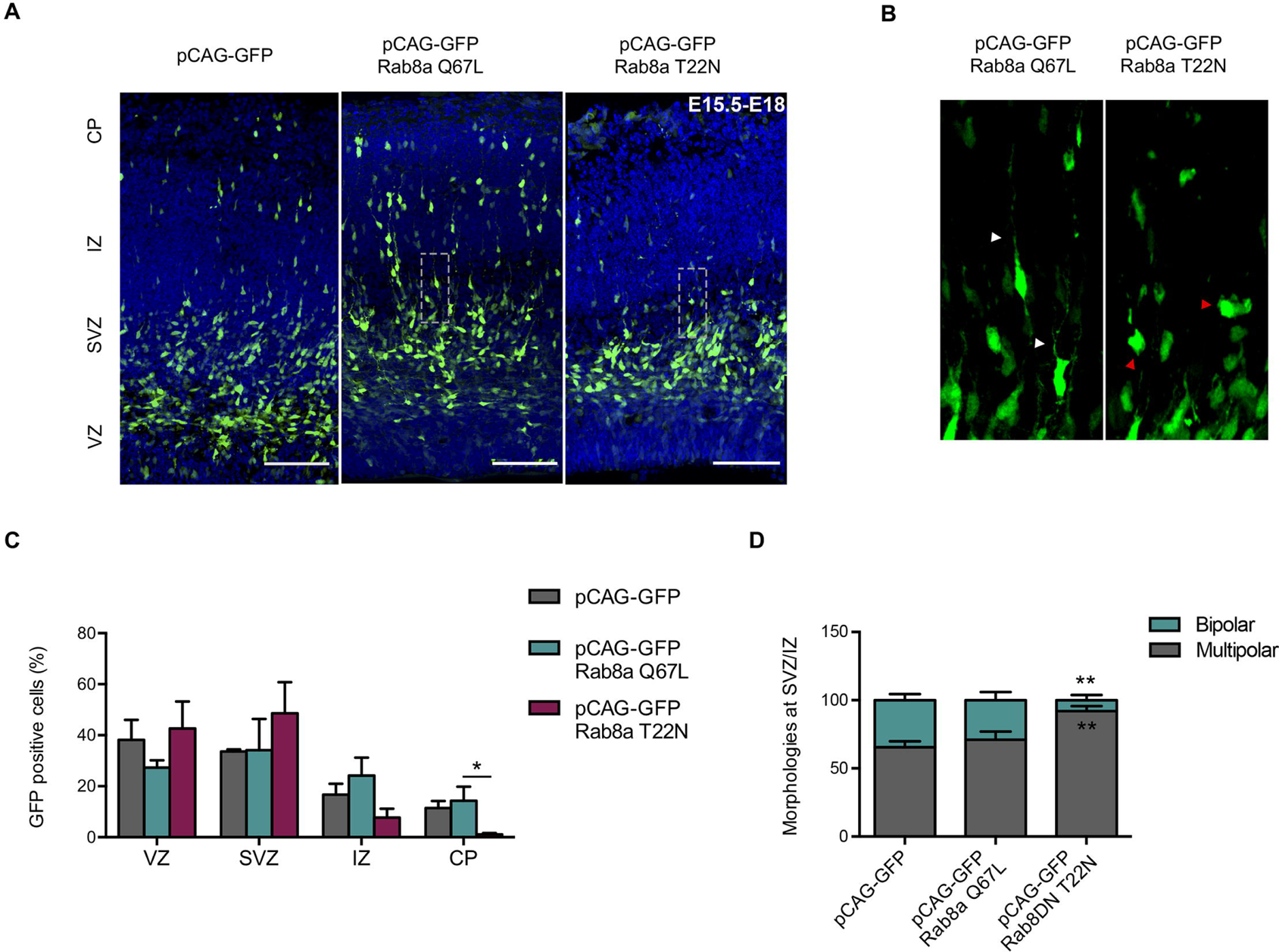
Rab8a regulates neuronal morphology of migrating neurons in the embryonic cortex. (a) Representative images of cortical sections of in utero-electroporated mice with pCAG-GFP, pCAG-GFP plus Rab8a Q67L or pCAG-GFP plus Rab8a T22N, at E15.5 and analyzed at E18. GFP positive cells were located in the cortical plate (CP), intermediate zone (IZ), subventricular zone (SVZ) and ventricular zone (VZ). **(b)** Representative image of a zoom between SVZ and IZ of panel (a). The black arrowhead shows normal neurons with a bipolar morphology, whilst the red arrowhead shows cells with a multipolar morphology. **(c)** Quantification of the distribution of GFP-positive cells in CP, IZ, SVZ and VZ. **(d)** Quantification of the morphology of GFP-positive cells between IZ and SVZ. Scale bars, 75 μm. Values represent mean ± SEM (n=4). *p<0.05 and **p<0.01 as compared to the corresponding control.

Neurons *en route* toward the apical surface of the cortex displayed a distinctive migration morphology characterized as bipolar, with a leading process that later developed into apical dendrites and a trailing process that is initially specified as the axon (Yogev and Shen, 2017). Therefore, we analyzed the presence and prevalence of multipolar and bipolar GFP-positive neurons in the SVZ-IZ area in control and Rab8a T22N or Rab8a Q67L mutants (Figure 7A-B, D). We found that control and Rab8a Q67L-expressing neurons display similar proportions of multipolar to bipolar morphologies. In contrast, neurons expressing Rab8a T22N, a DN mutation, were significantly more multipolar, reaching 92% of the neurons analyzed. Moreover, several Rab8a T22N-expressing neurons displayed a round morphology (Figure 7B, red arrowhead). Concomitantly, the proportion of bipolar neurons, that correspond to proper polarized migrating neurons, was significantly reduced. Therefore, our *in vivo* results indicate that Rab8 regulates neuronal polarity acquisition and migration, reinforcing the notion that Rab8a plays a critical role instructing axon determination and outgrowth.

## DISCUSSION

### Role of Tuba in neuronal polarity acquisition

In this report, we show that two small GTPases, Rab8 and Cdc42 are crucial molecular determinants for axon specification, and that they are molecularly-linked by Tuba. Tuba is a multidomain scaffolding protein that consists of four N-terminal SH3 domains that bind directly to dynamin1, an internal BAR (Bin, Amphiphysin, Rvs) domain, a GEF domain specific for the Rho-family GTPase Cdc42 (generating an intermediate activation) and two C-terminal SH3 domains. The extreme C-terminal SH3 domain directly binds to multiple actin-regulatory proteins, including N-WASP and Ena/VASP (Kovacs et al., 2006; Salazar et al., 2003). There are two isoforms of Tuba generated by alternative splicing, that differ by the presence of N-terminal SH3 domains, named TubaFL (∼180 kDa) and mTuba (∼75 kDa). TubaFL is an important factor in epithelial cell polarity. In 2D cultures, Tuba localizes at the apical tip of the tight junctions, and is required for the normal maturation and maintenance of adherens junctions. In this model, Tuba-mediated activation of Cdc42 leads to the activation of both the N-WASP and Par6–aPKC pathways (Otani et al., 2006). Additionally, Tuba is pivotal for the correct polarization and lumen formation in 3D MDCK spheroids, a model used to elucidate the mechanisms of epithelial morphogenesis and luminogenesis. Indeed, apical activation of Cdc42 depends on the coincidence between phosphatidylinositol 4,5 biphosphate (PIP2) patches at the plasma membrane and the presence of Tuba in the apical cytosol, allowing Par3–aPKC–Cdc42 complex activation (Bryant et al., 2010). However, its role in neuronal polarity has not been addressed previously. Similarly to epithelial cells, in developing neurites, activation of the Par6-Par3–aPKC-Cdc42 cascade is required for axon growth (Lalli, 2009), but the identity of the GEF involved in Cdc42 activation had not been described until now. Our results indicate that Tuba connects Rab8 and Cdc42 activities which are essential to define axon identity. Tuba enrichment at the axonal growth cone tip, recapitulates the behavior of other proteins involved in neuronal polarity; i.e. before axon specification, Par3 is localized in the cell body and at the tips of all nascent processes (Nishimura et al., 2005; Schwamborn and Puschel, 2004; Shi et al., 2003), whereas in stage 3, Par3 is lost from minor neurites and becomes selectively-expressed in the axon and developing growth cone (Shi et al., 2003). The spatial and temporal expression of Par6 during axon specification is similar to that of Par3, being confined to the cell body and the axon at stage 3 (Schwamborn and Puschel, 2004; Shi et al., 2003). The relevance of Tuba in axon specification is reinforced by loss- and gain-of function approaches that highlight antagonistic effects over neuronal polarity acquisition. Consistently, loss- and gain-of-function for Cdc42 lead to similar results (Garvalov et al., 2007; Govek et al., 2018; Lopez Tobon et al., 2018; Schwamborn and Puschel, 2004). A truncated version of Tuba, called mTuba had no effect on neuronal polarity. Unlike mTuba, TubaFL can bind dynamin (Kovacs et al., 2006) and GM130, controlling Cdc42 activation at the Golgi apparatus (Kodani et al., 2009).

### The small GTPase Rab8, a relevant factor in neuronal polarity

The secretory pathway is the primary source for newly-synthesized membranes and is essential for axon spreading and growth. Rab8 was originally described as a trafficking regulator between the TGN and the plasma membrane and its overexpression induces the formation of long cell surface protrusions in a variety of cells (Hattula and Peranen, 2000; Huber et al., 1995; Peranen et al., 1996). A role for Rab8 in neurite elongation was proposed earlier (Huber et al., 1995); however its role as an axon determinant is novel. There are two Rab8 isoforms, Rab8a and Rab8b, which are encoded by two different genes. A recent report using high throughput screening shows that both isoforms are present in neurons, although Rab8a is expressed at higher levels (Sharma et al., 2015). We analysed the effect of both Rab8 isoforms, and concluded that only neurons overexpressing Rab8a CA have supernumerary axons, whereas neurons overexpressing Rab8a DN or Rab8b DN have a greater number of neurons arrested in stage 2, without axon generation. In contrast, a previous study in primary mouse cortical neurons, reports that overexpression of Rab8a CA generates longer axons than control neurons, whereas neurons overexpressing Rab8a DN does not have differences in axonal length compared to control neurons. Those authors suggested a coupling between Rab8 and Rab11 functions, likely at the recycling endosomes (Furusawa et al., 2017). A role for Rab8 in axon specification is also supported by *in vivo* experiments showing that reducing Rab8a activity in the brain significantly increases the proportion of multipolar neurons in the cortex. Similar phenotypes were found by targeting other molecular components involved in neuronal polarity acquisition such as IGF-1R (Nieto Guil et al., 2017) and Par3 (Funahashi et al., 2013).

Several lines of evidence suggest that key processes determining neuronal morphology, such as axon specification, elongation and branching, could be dissected at a molecular level, based on their dependence on Rab-specific vesicular trafficking regulation (Villarroel-Campos et al., 2016a; Villarroel-Campos et al., 2014). The Early/Late Endosome Rab GTPases: Rab5, Rab7, Rab21 and Rab22 have been mainly associated with the regulation of neurite outgrowth. The Recycling Endosome-Associated Rab GTPases: Rab4, Rab11, Rab35 have been linked with the regulation of axonal elongation. Finally, TGN-Related Rab GTPases: Rab10 and Rab33 have been reported to regulate axonal elongation and axonal specification, respectively (Villarroel-Campos et al., 2016a; Villarroel-Campos et al., 2014). The axon determination during neuronal polarization is preceded by bulk cytoplasmic flow and polarized TGN-derived vesicle transport towards one specific neurite (Bradke and Dotti, 1997; Calderon de Anda et al., 2008). Thus, Rabs involved in traffic regulation from the Golgi apparatus could be relevant in axonal specification. Therefore, we envision the existence of two pools of Rab8, one in the recycling pathway that regulates axonal elongation, and a second pool that controls exocytic traffic from the TGN regulating axonal specification.

We also found a connection between Rab8a and actin dynamics. We observed that Rab8 activation induces a Tuba-mediated increase in Cdc42 activity, and that the effect of Rab8a overexpression on neuron morphology is dependent upon Tuba expression.

### A positive feedback loop between Rab8a and Cdc42 regulating neuronal polarization

Positive feedback circuits are crucial during neuronal polarization, contributing to stochastic selection of one of the minor neurites to become the axon, and simultaneously preventing the axonal identity of the other minor neurites. Examples of positive feedback during neuronal polarity acquisition are observed during the coupling of Ras, PI3K and Cdc42/Par3/Par6/Rac1 signalling pathways (Barnes and Polleux, 2009). We observed that Cdc42 fast cycling overexpression cannot overcome the phenotype induced in the Rab8a DN mutant. Concomitantly, Rab8a CA overexpression cannot reverse the phenotype induced by the Cdc42 fast cycling mutant (Cdc42F28L), suggesting a reciprocal regulation. Previous reports showed that Cdc42 also regulates Golgi organization and post-Golgi traffic (Friesland et al., 2013; Kage et al., 2017) emphasizing the importance of post-Golgi traffic and therefore a putative involvement of Rab8 in this process. Additionally, in mouse intestine, another highly polarized tissue, Cdc42 deficiency impaired both Rab8a vesicle trafficking to the midbody during cytokinesis and Rab8a activation and its subsequent association with effectors (Sakamori et al., 2012).

In conclusion, the molecular mechanism here described positions Tuba as the Cdc42 GEF involved in axon specification, and proposes Tuba as a linker that connects cytoskeleton dynamics and the directed vesicular traffic during neuronal polarization.

## Supporting information

Supplemental Figura 2

## Acknowledgements

This work was supported by CONICYT grants to CG-B under the Fondecyt (#1180419) and FONDAP (#15150012) programs. PU was supported by the Fondecyt (#3160630) Postdoctoral program. CG-B and CC were supported by the IBRO Prolab grant. We thank Dr. Michael Handford for language support.

---

**Supplementary Figure 1. Tuba knockdown efficiency in the neuroblastoma cell line.** Representative western blot and quantification of Tuba expression in N1E-115 neuroblastoma cell lines overexpressing sh-scrambled, sh-Tuba A, sh-Tuba B, sh-Tuba C or sh-Tuba D. Values represent mean ± SEM (n=3). *p<0.05 as compared to the corresponding control.

**Supplementary Figure 2. Rab8 knockdown efficiency in the neuroblastoma cell line. (a)** Representative western blot and quantification of Rab8 expression in B104 neuroblastoma cell lines overexpressing sh-scrambled, sh-Rab8a A, sh-Rab8a B, sh-Rab8a C or sh-Rab8a D. **(b)** Representative western blot and quantification of Rab8 in B104 neuroblastoma cell lines overexpressing sh-scramble, sh-Rab8b A, sh-Rab8b B, sh-Rab8b C or sh-Rab8b D. Values represent mean ± SEM (n=3). *p<0.05 as compared to the corresponding control

**Supplementary Figure 3. Neurons overexpressing Rab8a CA have several axonal initial segments (AIS).** Representative image of Rab8a CA-transfected neurons (grey) stained with anti-Ankyrin G (AIS marker, red) and anti-Map1b (neuronal marker, green) antibodies at 14 DIV. Arrowheads indicate the AIS. Scale bars, 50 μm

