## Supplementary figures and images for "Tuba activates Cdc42 during neuronal polarization downstream of the small GTPase Rab8a"

### Supplemental Figura 2

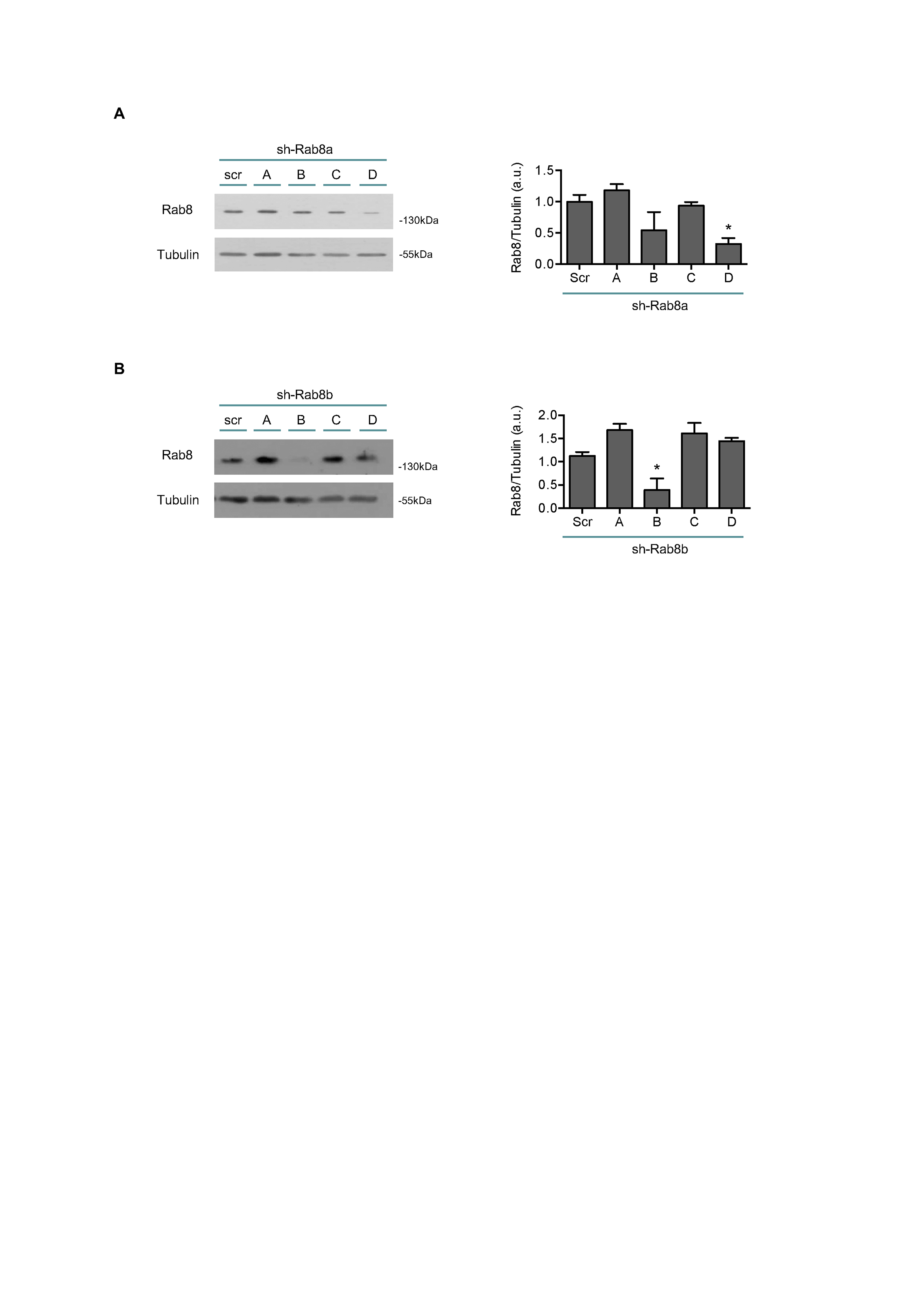

### Supplemental Figure 1

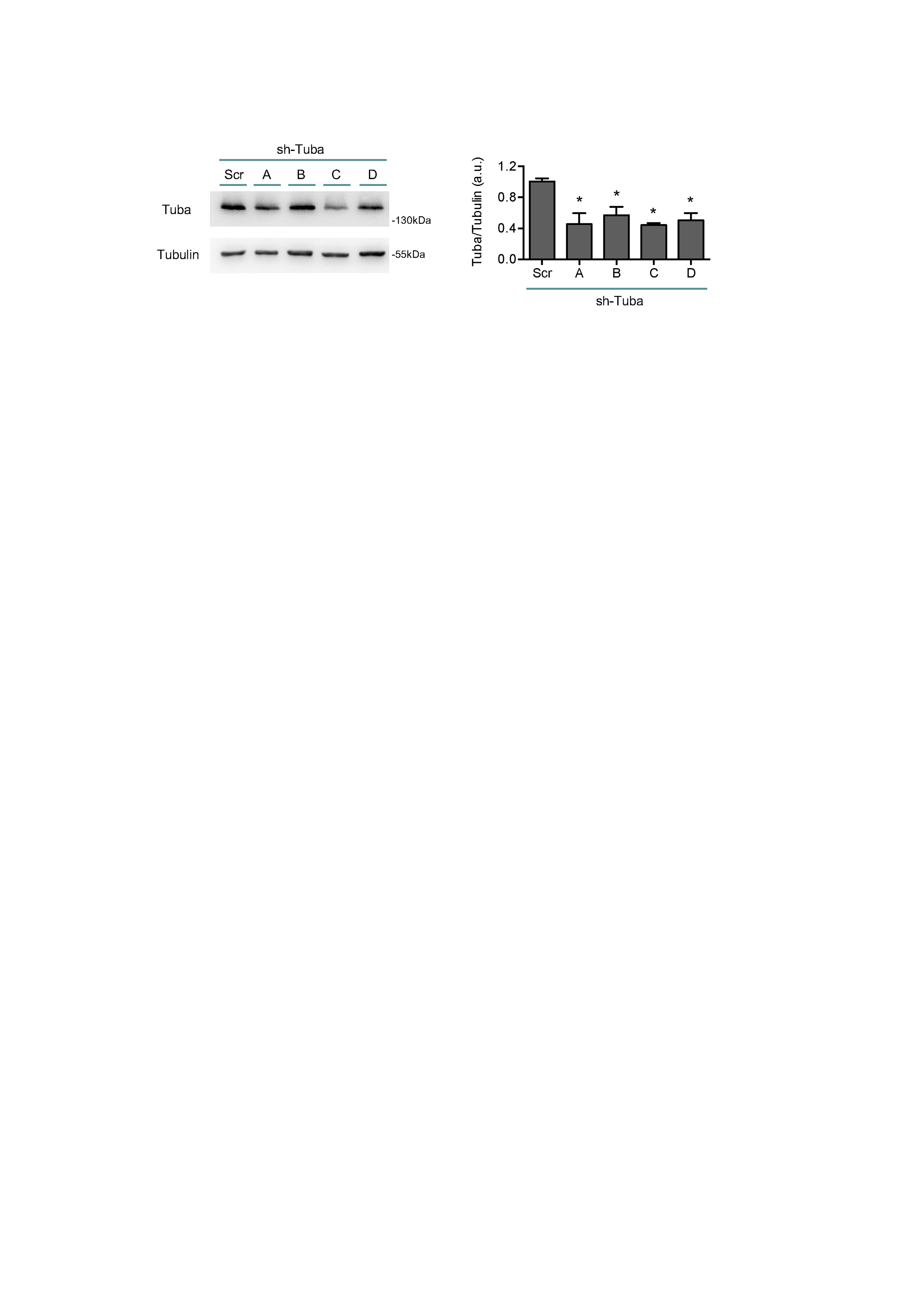

### Supplemental Figure 3

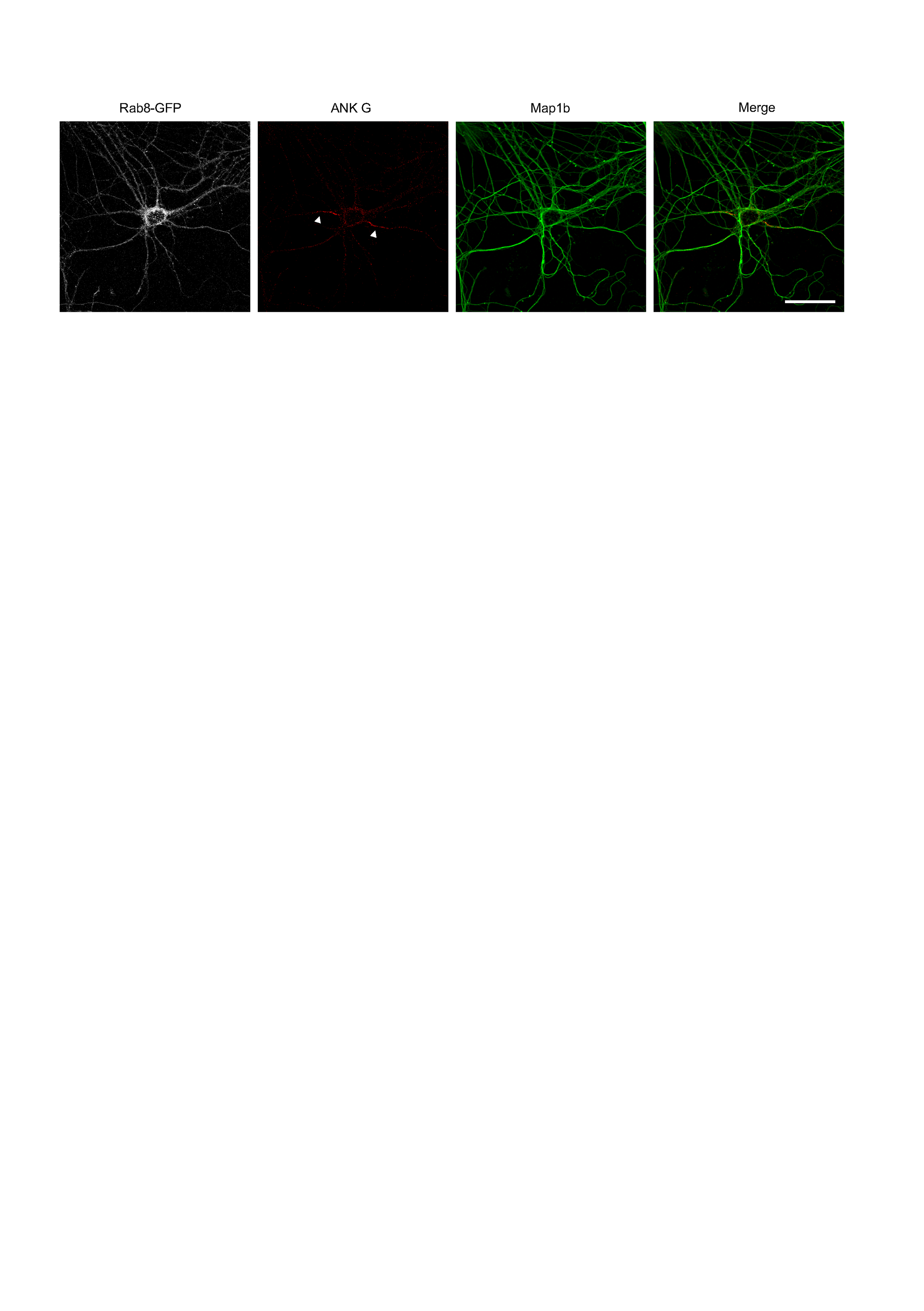
